# The lateral habenula integrates age and experience to promote social transitions in developing rats

**DOI:** 10.1101/2024.01.12.575446

**Authors:** Dana Cobb-Lewis, Anne George, Shannon Hu, Katherine Packard, Mingyuan Song, Oliver Nguyen-Lopez, Emily Tesone, Jhanay Rowden, Julie Wang, Maya Opendak

**Affiliations:** Kennedy Krieger Institute, Baltimore MD USA 21205; Solomon H. Snyder Department of Neuroscience, Johns Hopkins University School of Medicine, Baltimore MD USA 21205

**Keywords:** habenula, adversity, mPFC, development, social behavior

## Abstract

Social behavior deficits are an early-emerging marker of psychopathology and are linked with early caregiving quality. However, the infant neural substrates linking early care to social development are poorly understood. Here, we focused on the infant lateral habenula (LHb), a highly-conserved brain region at the nexus between forebrain and monoaminergic circuits. Despite its consistent links to adult psychopathology, this brain region has been understudied in development when the brain is most vulnerable to environmental impacts. In a task combining social and threat cues, suppressing LHb principal neurons had opposing effects in infants versus juveniles, suggesting the LHb promotes a developmental switch in social approach behavior under threat. We observed that early caregiving adversity (ECA) disrupts typical growth curves of LHb baseline structure and function, including volume, firing patterns, neuromodulatory receptor expression, and functional connectivity with cortical regions. Further, we observed that suppressing cortical projections to the LHb rescued social approach deficits following ECA, identifying this microcircuit as a substrate for disrupted social behavior. Together, these results identify immediate biomarkers of ECA in the LHb and highlight this region as a site of early social processing and behavior control.

## Introduction

Social behavior is critical for access to key resources such as food, protection, and receptive mates. For altricial species, the caregiver is the primary provider of such key resources, which requires infants to have a strong bias towards approaching the caregiver ^1,2^. Therefore, in early life, circuits that promote social approach behavior are engaged, while those which support avoidance are inactive or suppressed by the presence of the caregiver ^3–7^. As animals mature and become more independent, neural circuits shift to promote a balance of approach and avoidance behaviors to support changing social demands ^8–11^.

Disrupted social behavior is a core symptom of many psychiatric conditions, including major depressive disorder (MDD), autism, and anxiety ^12–15^. Early life adversity, especially in the context of a caregiver, has been identified as a significant risk factor for development of these conditions ^7,16–23^. In fact, nearly one-third of adult mental health conditions are associated with exposure to trauma early in life ^24^. However, the mechanisms underlying the correlational links between early caregiving adversity (ECA) and adult outcomes remain elusive. Understanding the neurobiological processes linking ECA and psychopathology is critical to developing effective neural interventions to target social deficits.

The lateral habenula (LHb) is considered the “anti-reward” region of the brain, as it responds to aversive cues and the absence of expected rewards ^25–28^. Through this signaling, the LHb serves as a functional locus for behavioral flexibility in tasks with changing reward and punishment contingencies ^29–31^, such as when evaluating social stimuli as safe or threatening. In fact, the LHb is implicated in a variety of adult social behaviors, including maternal behavior ^32–36^, social play ^37^, social isolation ^37,38^, and aggression ^39^. Activation of the LHb inhibits adult social behavior ^40^. Additionally, the LHb modulates social behavior via connections to forebrain regions and monoaminergic circuits (for review see ^41^). For example, we previously showed that, downstream of the LHb, the dopaminergic projection to the basolateral amygdala inhibits social approach in infants ^8^. In adults, the medial prefrontal cortex (mPFC) projection to the LHb conveys stress information ^42^ and suppresses social approach ^40^. This suggests that, in adults, the LHb sits at the interface of the mPFC and midbrain to modulate the influence of the cortex on dopaminergic encoding of reward value during social interactions. However, the role of the LHb in infant social behavior has been underexplored.

LHb dysfunction is consistently implicated in adult psychiatric conditions with their roots in development, including MDD ^43–54^. However, despite strong evidence of the role of the LHb in psychopathology, how ECA alters the LHb in the developing brain to initiate the pathway to the pathology remains unknown. Here, we use structural and functional assays to assess the development of the LHb in both typical development and in the context of ECA. We focus our assessments on the infant to juvenile developmental window, a transitional period marked by changing environmental demands and increased vulnerability to environmental impacts. We show that (1) ECA blunts growth of the LHb, (2) ECA increases bursting activity of LHb neurons, (3) ECA alters developmental expression of neuromodulatory receptors in the LHb, (4) there is a developmental switch in the role of the LHb during a task involving social behavior flexibility, and (5) impaired infant social approach following ECA is rescued by suppressing mPFC projections to the LHb. Altogether, these results identify the LHb as a substrate for early social behavior flexibility and a locus of dysfunction in the corticolimbic circuit following stress.

## Materials and Methods

### Subjects

Male and female Long Evans rats were born and bred at Kennedy Krieger Institute (originally from Envigo). They were housed in polypropylene cages (23L x 20H x 46W cm) with corncob bedding in a 72 ± 5°F temperature environment on a normal 12h light/dark cycle. Day of birth was considered PN0 and litters were culled to 12 pups (6 males, 6 females) on postnatal day (PN) 1. Food and water were available *ad libitum.* Pups were separated from their mother only for the duration of the surgery and/or behavior sessions (maximum 1h). Weaning occurred at PN21. All procedures were approved by the Johns Hopkins University Animal Care and Use Committee.

### Scarcity-adversity model of low bedding (SAM-LB)

Early life care was disrupted using a scarcity-adversity model of low-bedding (SAM-LB) that has been well established in previous literature ^55–59^. All pups received normal rearing from PN0-PN8, then control or SAM-LB rearing from PN8-12. These ages were selected based on previous research from our laboratory and others that suggests experiencing adversity during this age range produces enduring neurobehavioral effects across development ^55,57,60,61^.

Experimental litters were switched to wood shaving bedding no later than PN5. In the SAM-LB condition, bedding resources were restricted to 100 mL, leading to more frequent rough handling of pups (Table 1). Litters were maintained on a solid floor and the cage was not cleaned over the course of the treatment, consistent with previous studies using this model ^8^. Makeshift nests created by the mother using food or feces were removed as needed. Maternal behaviors were validated from 1-hour videos filmed on three out of the five treatment days for both control and SAM-LB reared pups. Maternal behavior changes over the course of the treatment are shown in Table 1.

**Table 1.** Frequency of maternal behaviors across litters in each rearing condition.

|  | Control | SAM-LB Adversity |
| --- | --- | --- |
| Dam in nest (% observations) | 86.67 ± 6.24 | 86.67 ± 9.72 |
| Nursing (% observations) | 33.33 ± 5.27 | 43.33 ± 19.44 |
| Grooming (% observations) | 66.67 ± 9.13 | 83.33 ± 12.91 |
| Rough handling<br>(% observations) | 3.33 ± 3.33 | 73.33 ± 16.33*<br>*t*(4)=-4.199, *p*=0.014 |
| Pup weight (g) | 24.47 ± 1.12 | 24.03 ± 1.87 |

### Habenula volume assay

To obtain volume measurements for the medial and lateral habenulae, separate cohorts of pups received SAM-LB or control rearing from PN8-12 and were perfused with ice cold 4% PFA at PN14 or PN28 and brains stored until sectioning (40 µm) on a microtome. All litters used for the volume study had 10-12 pups and average litter size was constant across groups. Separate litters were used for volume assessments at each age. Sections were stained with DAPI and area measurements taken using ImageJ software (NIH) by counting the number of slices containing the habenula and manually tracing the area of the largest LHb and medial habenula (MHb) as defined by Paxinos and Watson ^62^. The volume of each region was calculated using Cavalieri’s principle using the formula for a cone ^7,63^.

### *In vivo* electrophysiology

Male and female rat pups (PN12-18 and PN26-28) were anesthetized with urethane (1.8 g/kg, i.p.) and placed in an adult stereotaxic apparatus modified for use with infants. A heating pad (Pain Care Labs, Atlanta, GA) was used to maintain body temperature. Moveable array 16-channel microdrives (Innovative Neurophysiology, Durham, NC) were stereotaxically positioned unilaterally (hemisphere counterbalanced across subjects) within the LHb (P18: -2.8 AP, ± 0.7 ML, -3.6 to -4.6 DV; P28: -3.2 AP, ± 0.8 ML, -3.8 to -4.8 DV) and used to record multi-unit activity. A single channel without detectable units was used as a reference electrode. Signals were amplified and digitized with a SmartBox Pro (NeuroNexus, Ann Arbor, MI), and acquired using Allego Radiens software (NeuroNexus, Ann Arbor, MI). Data were sampled at 30 kHz and filtered (300-5000 Hz bandpass; 60 Hz notch). Microdrives were painted with Vybrant DiI labeling solution (Invitrogen, Waltham, MA) prior to implantation to verify recording position *post hoc*.

### Spike Sorting & Analysis

Spikes were sorted and classified offline using Spike2 software (CED, Cambridge, ENG). Single units were identified by threshold crossings and principal component analysis (PCA). Spikes with interspike intervals less than the refractory period (2 ms) were excluded. Firing parameters, including firing rate, bursting, and coefficient of variation (CV) were analyzed offline using Spike2 software(CED, Cambridge, ENG). The CV was calculated as a ratio of the standard deviation and mean of the interspike interval (ISI) of each neuron.

Burst events were detected using the Spike2 script, *Bursts*. Bursts were defined using previously established criteria of ≤ 80 ms ISI to signal the onset of a burst, ≥ 160 ms to signal the end of a burst, and a minimum of 2 spikes per burst ^64^. Cells were separated into firing pattern clusters (bursting or non-bursting) using two parameters: percent spikes fired as bursts (%SFB) and the CV of the ISI. ISI histograms were also generated to aid in identification of firing patterns.

### Lateral habenula protein expression assay

At PN14 or PN28, the habenula was rapidly dissected on ice following decapitation. Tissue was then immediately frozen on a metal plate cooled by dry ice, transferred to Eppendorf tubes, and stored at -80°C prior to homogenization and protein quantification. Tissue was homogenized in 300 μl of RIPA lysis buffer (Santa Cruz Biotechnology, Dallas, TX), containing PMSF (200 mM), protease inhibitor cocktail, and sodium orthovanadate (100 mM), using a sonicator (Qsonica, Newton, CT) at level 30 for five seconds. Homogenized samples were incubated on a rocker for one hour at 4°C and then centrifuged at 10,000 rpm for 10 minutes at 4° C. The resulting supernatant was collected and kept on ice. The Pierce bicinchoninic acid assay (BCA; ThermoFisher, Rockford, IL) was used to determine protein concentration for each sample. Samples were reduced with 6X Laemmli sample buffer (Thermo Scientific, Waltham, MA) equivalent to 1/6 of the total sample volume, boiled for five minutes, and stored at -20°C prior to immunoblotting. 30 µg of protein from each sample was loaded into Bolt 4-12% Bis-Tris Plus gels (Invitrogen, Waltham, MA). Each gel contained at least one control PN14 sample as a comparison value for calculating fold change from this predetermined baseline. Samples were transferred onto nitrocellulose membranes (Invitrogen, Waltham, MA) using the iBlot Dry Blotting System (Life Technologies, Carlsbad, CA) for seven minutes. Membranes were then blocked in 5% BSA for 1 hour at room temperature, and then incubated in primary antibodies against the first protein of interest and tubulin (Abcam, Cambridge, UK,; 1:5000) overnight at 4°C. Proteins of interest included Ca2+/calmodulin-dependent protein kinase II β (CaMKIIβ; Abcam, Cambridge, UK,; 1:1000), GABA_B_ receptor (Abcam, Cambridge, UK,; 1:1000), estrogen receptor 1 (ESR1; Abcam, Cambridge, UK,; 1:1000), serotonin receptor 2C (5-HTR2C; Invitrogen, Waltham, MA; 1:1000). Membranes were washed three times in 0.1% Tween-20 in 1X TBS (TBST) for five minutes each and incubated in 800 nm and 700 nm dye-conjugated secondary antibodies (Li-Cor; 1:10,000) for 1 hour at room temperature for multi-channel imaging on a Li-Cor reader.

Following imaging, membranes were incubated in stripping buffer (Thermo Scientific, Waltham, MA) for 15 minutes, washed three times in TBST for five minutes each, incubated in primary antibody against a second protein of interest overnight, and incubated in 800 nm or 700 nm dye-conjugated secondary antibody for 1 hour at room temperature. Blots were stripped one time at maximum. Bands were analyzed in ImageJ (NIH) and normalized to tubulin levels. Fold change from control, PN14 control-reared animals, was calculated and visualized on graphs.

### Metabolic imaging via 2-DG autoradiography

At PN18 or PN28, pups were injected (20μCi/100g, s.c.) with 14C 2-deoxyglucose (2-DG) and placed in a Plexiglass beaker. Pups at each age then underwent a 10-minute habituation period, followed by eight 0.5 mA shock to the tail (4 min ITI) either alone or in the presence of an anesthetized dam, according to previously published protocols ^8,65^. 45 minutes after the procedure ended, pups were perfused and brains were removed for analysis of 2-DG autoradiography using ImageJ (NIH), following previous protocols ^66–68^.

For functional connectivity modeling, bivariate correlation matrices were created by computing ratios of mean 2-DG uptake for all pairwise combinations of brain regions. For quantitative analyses, each group’s correlation matrices were transformed into z-scores and group differences between modules were analyzed by ANOVA and *post hoc* Fisher’s LSD.

### General infant surgery procedure

Pups were removed from the nest without disturbing the mother and placed partially on a surgical heating pad to permit thermoregulation. No more than 6 out of 12 pups from a litter were used to minimize stress to the mother. Before surgery, pups were anesthetized by inhalation with isoflurane and placed in an adult stereotaxic apparatus modified for use with infants. During aseptic survival surgery, visual inspection of skin color, motor tone and respiration were continuously performed to monitor for signs of distress or waking. Pups were kept on a heat pad throughout the surgery. Following craniotomy, an injector cannula was lowered into the LHb (-1.8 AP, ± 0.5-0.6 ML, -2.5 DV) and/ or the mPFC (1.2 AP, ± 0.5-0.6 ML, -1 DV) using coordinates relative to Bregma verified from an infant rat atlas ^69^. The cannula was lowered 0.1mm beyond the DV coordinate, left for one minute, and raised to the DV coordinate before infusion. Viruses (UNC Vector Core) were infused at 0.1µL/min for a total volume of 0.3µL per side. The cannula was left in place for one minute following the end of infusion to ensure viral spread.

For LHb chemogenetic experiments, PN3-5 pups were bilaterally transduced with AAV9-hSyn-hM4D(Gi)-mCherry or AAV5-hSyn-mCherry into the LHb. For mPFC-Hb chemogenetic experiments, at PN3-5 pups were bilaterally transduced with Cre-dependent AAV9-hSyn-DIO-hM4D(Gi)-mCherry or AAV9-hSyn-DIO-mCherry in the LHb and AAVrg-hSyn-Cre-P2A-dTomato in the mPFC using two cannulae simultaneously. This Cre-dependent approach specifically targets neurons projecting from the mPFC to the LHb. For activity-dependent tagging experiments, PN3-5 pups were bilaterally transduced with a 1:1 mixture of AAV5/cFos-ERT2-Cre-ERT2-PEST-P2A and AAV9-hSyn-DIO-mCherry in the LHb. Some animals were surgerized at PN8-9 to control for virus expression time when comparing manipulations across age.

The incision was stitched with 2-3 sutures. Following a 30 min recovery and regaining of righting reflex, pups were returned to the nest until experimental procedures. One day of recovery is sufficient for pups up to weaning age ^8^. Pups were anesthetized for <20 min and were not away from the mother >60 min ^55,60,70–73^.

#### Ensuring pups thrive after surgery

Following all surgical procedures, pups and mothers were inspected after pups were returned to the nest to ensure pups were not rejected and were able to thrive (i.e. typical behavior and presence of milk band, based on published procedures from the Sullivan lab ^7,55,70,74–79^). Before being returned to the nest, experimenters ensured pups were fully recovered from anesthesia and were mobile, responsive to tactile stimulation, and behaving indistinguishably from untreated pups. All removals and returns to the nest took less than one minute and were performed by trained experimenters to ensure minimal disruption for the dam. Dam behavior was inspected for the following atypical behaviors: dragging pups, rough handling, and rejecting experimental pups. The animals were euthanized if they presented failure to gain weight or failure to respond normally during the preparation for the experiment. For example, vocalizations during handling indicate the animal did not recover well after surgery. Over the course of the study, <10 pups were euthanized for these reasons.

### Social behavior testing

Cohorts were tested at PN17 and PN27 for social behavior using an adaptation of the Crawley 3-chamber social test apparatus (3CT) ^14^. Animals were habituated to the chamber for 5 minutes prior to the social behavior test. Then a novel peer (same age as stimulus) was placed into one of two perforated, transparent Plexiglas boxes (6x6x6 in) in each of the outside quadrants (location of social stimulus counterbalanced) for 12-minutes. Time spent in each chamber, the duration of close proximity to the box containing the novel peer, and activity level were recorded and automatically analyzed using the Noldus Ethovision XT system (Leesburg, VA). Boxes were cleaned with 70% ethanol between trials. Videos were validated by handscoring using BORIS software ^80^. Peers were observed for stress responses, including freezing, hyperactivity and hypervigilance. We observed no evidence of stress in peers, consistent with previous studies showing that neither stimuli nor experimental animals emit ultrasonic vocalizations in the three-chamber test ^81^ and habituation to the chamber before testing prevents additional stress effects ^75,82^. At PN18 and PN28, animals underwent the same social behavior test with the addition of ambient predator odor (5 µl fox urine (Leg Up Enterprises, Lovell, ME) placed on a Kimwipe in one of the top corners of the box). All animals were habituated to the chamber with the presence of the predator odor for 5 minutes prior to the social behavior test. Intraperitoneal (i.p.) injections (3 mg/kg) of clozapine-N-oxide (CNO dihydrochloride; HelloBio, Princeton, NJ) were given to all animals 30 minutes before each test session. After the juvenile social behavior test with predator odor, animals were independently housed for an hour, before being sacrificed and perfused.

### Activity-dependent neural tagging

Population tagging was performed via a single intraperitoneal (i.p.) injection of either 0.9% saline or 40 mg/kg 4-hydroxytamoxifen (4-OHT; Sigma, St. Louis, MO, Cat# H627) immediately following sociability testing in the presence of predator odor on PN16-17 ^83–85^. 4-OHT was prepared fresh the day of use. Briefly, 4-OHT was dissolved and diluted to a concentration of 4 mg/mL in saline with 2% Tween-80 and 5% DMSO. Following 4-OHT injection, pups were returned to their homecage with littermates and dam. This age range was selected to maximize recombination time while closely matching the chemogenetic timeline. Immediately following behavior testing at PN27-28, animals were independently housed for one hour, before being sacrificed and perfused. We also utilized homecage controls that received a single i.p. injection of 4-OHT within the homecage on PN16-17 and were sacrificed and perfused directly from the homecage on PN27-28. Tissue was processed for mCherry and c-Fos.

### Immunohistochemistry

Rats were anesthetized and transcardially perfused with ice cold 4% paraformaldehyde (PFA) in PBS (pH 7.4). Brains were postfixed in 4% PFA and then cryoprotected in 30% sucrose solution. 40 µm coronal slices were sectioned on the microtome and then subsequently were stored at 4°C in PBS with 0.01% sodium azide until processing. Sections were washed in PBS and then incubated for 20 minutes at room temperature in 3% normal horse serum. A solution of primary antibodies (1:500 rabbit anti-mCherry, Invitrogen #PA5-34974 or Abcam, Cambridge, UK, Cat #167453; and 1:500 guinea pig anti-cFos, Synaptic Systems, Göttingen, GER, Cat#226 308) was diluted in 3% normal goat serum and 0.05% Triton-X solution and then applied overnight at room temperature. Sections were then washed in PBS and a solution of fluorophore-conjugated secondary antibodies (1:300 horse anti-rabbit 594, Vector, Olean, NY, Cat#DI-1094 and 1:200 goat-anti guinea pig 488, Abcam, Cambridge, UK, Cat#150185 or Invitrogen, Waltham, MA, Cat#A11073) were applied at room temperature for 30 minutes. Finally, sections were washed in PBS and mounted on slides with DAPI Fluoromount-G (Southern Biotech, Birmingham, AL, Cat#0100-20). To quantify activity-dependent tagging, LHb sections were imaged for mCherry and Fos and overlap manually counted.

### Statistical analysis

Data were analyzed using GraphPad Prism software (version 9.0) with Student’s t-tests, one-, two-, and three-way analysis of variance (ANOVA) and *post hoc* Sidak-corrected comparisons. Data in graphs are expressed as mean ±SEM. Differences with *p*<0.05 considered statistically significant.

## Results

### Experiment 1

#### Development and rearing impact habenular volume and protein expression

As alterations to habenular volume are associated with psychiatric disorders ^50,86,87^, we sought to understand how habenular volume changes across development and following ECA. To characterize structural impacts of rearing and age, we assessed volume of the habenula in the infant (PN14) and juvenile (PN28) rodents (Figure 1A). We did not observe an effect of lateralization, therefore data were collapsed across hemispheres in both the medial (3-way ANOVA, age x rearing x hemisphere, *F*_(1,48)_=0.038, *p*=0.847) and lateral (3-way ANOVA, age x rearing x hemisphere, *F*_(1,48)_=0.866, *p*=0.357) habenulae. In the medial habenula (MHb), we observed a main effect of age (Figure 1B; 2-way ANOVA, *F*_(1,24)_=21.59, *p*<0.001)*. Post hoc* comparisons revealed age-related increases in both control-reared (*p*=0.007) and SAM-LB-reared pups (*p*=0.005).

**Figure 1.**
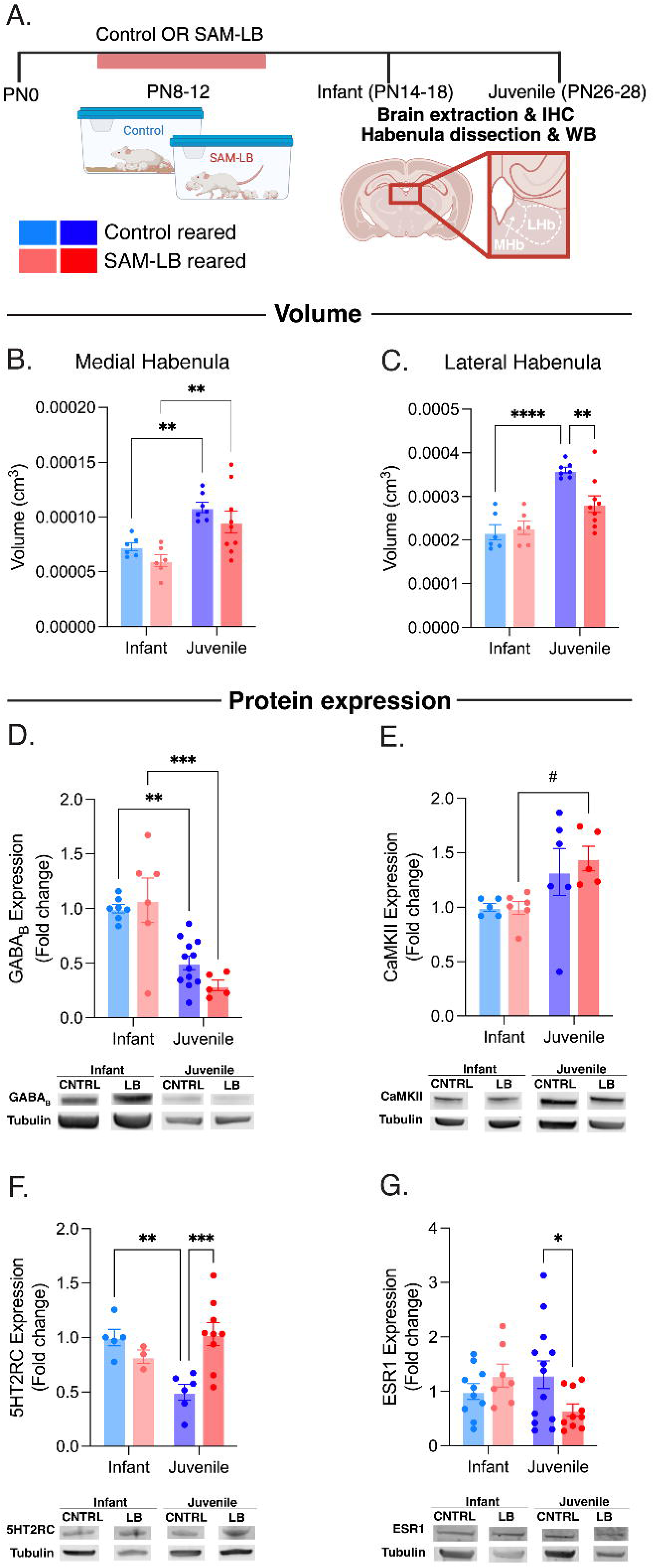
Development and rearing impact habenular volume and protein expression. (A) Schematic illustrating experimental timeline. (B) Average volume of the medial habenula. (C) Average volume of the lateral habenula. (D) Relative expression of GABA_B_ receptors in the lateral habenula with example blots. (E) Relative expression of CaMKIIβ receptors in the lateral habenula with example blots. (F) Relative expression of 5HT2C receptors in the lateral habenula with example blots. (H) Relative expression of ESR1 receptors in the lateral habenula with example blots. All data reported as mean ± SEM. \**p*<0.05, \*\**p*<0.01, \*\*\**p*<0.001, ^#^*p*<0.1

By contrast, in the LHb we observed an impact of adversity on volume. We observed a significant interaction between age and rearing (Figure 1C; 2-way ANOVA, *F*_(1,24)_=7.151, *p*=0.013) and a main effect of age (*F*_(1,24)_=35.49, *p*<0.0001). *Post hoc* comparisons revealed that the LHb was larger in juveniles compared to infants in control-reared pups (*p*<0.001). However, these age-related increases were not observed in SAM-LB-reared pups (*p*=0.074) such that LHb volume was smaller in juvenile SAM-LB-reared pups compared to control-reared pups of the same age (*p*=0.004). Together, these results suggest that, in typical development, both the lateral and medial habenulae increase in size from infancy to the juvenile period. However, ECA perturbs this growth curve.

To gain a more detailed understanding of LHb functional ontogeny, we next assessed impacts of development and rearing on protein expression in this region. Calcium calmodulin-dependent protein kinase II beta (CaMKIIβ), a protein expressed by excitatory neurons, plays a critical role in shaping the architecture and function of the developing nervous system, including dendritic morphogenesis and synaptic plasticity ^88,89^. GABA_B_ receptors similarly exert neurotrophic influence, while also contributing to network excitatory/inhibitory balance ^90,91^. Therefore, we wanted to explore how expression of these receptors change across development and following ECA.

We observed that expression of GABA_B_ receptors in the LHb decreases from infancy to the juvenile period (Figure 1D; 2-way ANOVA, main effect of age: *F*_(1,26)_=38.28, *p*< 0.001). *Post hoc* comparisons revealed this was statistically significant in control-reared (*p*=0.001) and SAM-LB-reared pups (*p*<0.001). Within age, groups did not differ based on rearing (all comparisons *p*>0.05). CaMKIIβ expression in the LHb shows the opposite developmental trend, increasing from infancy to the juvenile period (Figure 1E; 2-way ANOVA, main effect of age: *F*_(1,18)_=8.302, *p*=0.01; *post hoc* comparison *p*=0.057).

While the LHb is almost entirely glutamatergic ^92^, its activity is modulated by inputs from ventral tegmental area, dorsal raphe nucleus, and lateral hypothalamus ^93–96^, among many other inputs. Additionally, neuromodulatory receptors (e.g., serotonin 5-HT2C and estrogen ESR receptors) in the LHb are associated with anxiety and depressive-like behavior in rodents ^39,97,98^. Therefore, we investigated how development and early life experience impact neuromodulatory receptor expression in the LHb.

Interestingly, we observed an interaction between age and rearing for 5HT2RC expression (Figure 1F; 2-way ANOVA, *F*_(1,19)_=11.03, *p*=0.004) and ESR1 expression (Figure 1G; 2-way ANOVA, *F*_(1,36)_=5.136, *p*=0.03) in the LHb. *Post hoc* comparisons showed a significant decrease in 5HT2RC expression from infancy to juvenile in control-reared pups (*p*=0.005), but elevated expression in SAM-LB-reared juvenile pups compared to age-matched controls (*p*=0.001). Additionally, *post hoc* comparisons revealed SAM-LB-reared juvenile pups showed decreased ESR1 expression compared to control-reared juvenile pups (*p*=0.043). Additionally, Together, these findings suggest that there is a developmental increase in baseline LHb excitatory signaling from the infant to juvenile period in both typical and adversity-rearing contexts. The observed changes to CaMKIIβ and GABA_B_ receptors may reflect synaptic pruning occurring across the infancy and juvenile periods. Importantly, we find that ECA alters expression of neuromodulatory receptors in the LHb, which could further alter the excitability of LHb neurons.

### Experiment 2

#### LHb firing rate and composition transitions from infant to juvenile period

Hyperactivity in the LHb has been linked with the development of psychiatric disorders such as depression ^43–48^. Because we saw changes to excitatory and neuromodulatory markers in the LHb (see Figure 1), we next assessed impacts of development and rearing on LHb activity using *in vivo* extracellular multi-unit recordings in infant and juvenile anesthetized pups (Figure 2A-C). We found that there is an increase in activity from infancy to the juvenile period (Figure 2D; 2-way ANOVA, age x rearing, main effect of age: *F*_(1,143)_=10.90, *p*=0.001) in both control (*p*=0.044) and SAM-LB reared (*p*=0.039) pups. Interestingly, however, we also observed a reduction in the number of spontaneously active habenular neurons from infancy to the juvenile period (Figure 2E; 2-way ANOVA, main effect of age: *F*_(1,14)_=18.78, *p*=0.001) in both control-reared (*p*=0.018) and SAM-LB reared (*p*=0.015) pups. This observed developmental increase in excitability seems to be due to an increase in bursting activity, as the number of bursts per minute increased from infancy to the juvenile period (Figure 2F; 2-way ANOVA, main effect of age: *F*_(1,75)_=21.15, *p*<0.0001) in both groups (control, *p*=0.004 and SAM-LB, *p*=0.003). These data suggest a transition to more adult-like firing rates and patterns of firing across the infancy to juvenile period to include a mixture of bursting, silent, and tonically firing cells ^99,100^.

**Figure 2.**
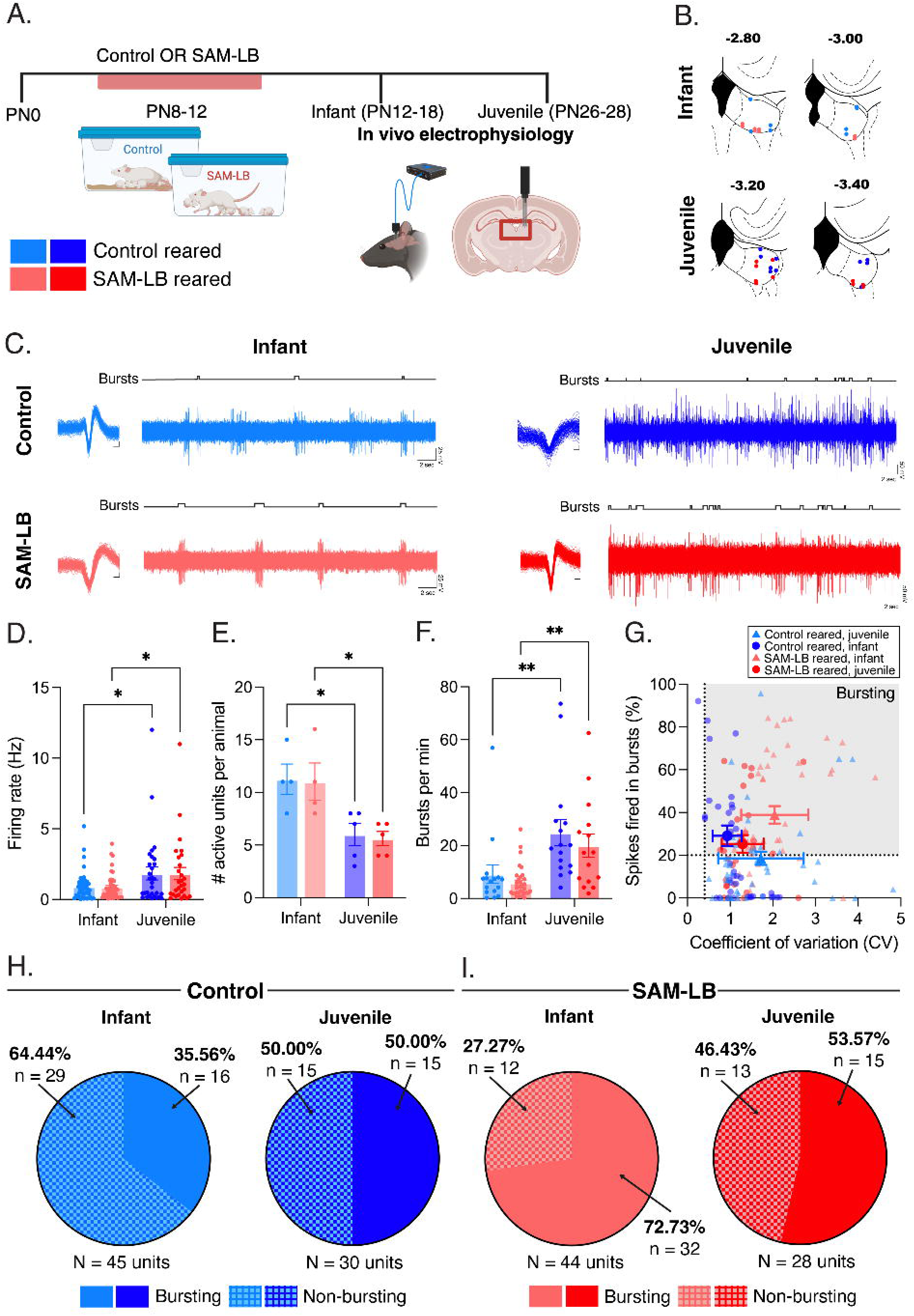
LHb firing rate and composition transitions from infant to juvenile period. (A) Schematic illustrating experimental timeline. (B) Map of recording locations in infant control (light blue), juvenile control (dark blue), infant SAM-LB (light red), and juvenile SAM-LB (dark red). (C) Representative extracellular *in vivo* recordings of spontaneous firing activity in LHb neurons. (D) Average spontaneous firing rate. (E) Average number of spontaneously active units per animal. (F) Average number of bursts per minute. (G) Scatterplot showing the coefficient of variation of the interspike interval (mean/SD) and percent of spikes fired in bursts (%SFB) of recorded LHb neurons. Line at SFB 20% and 0.4 CV represents the threshold for neurons classified as bursting. (H) Percent units classified as bursting (solid) and non-bursting (hatched) in infant (light blue) and juvenile (dark blue) control reared pups. (I) Percent units classified as bursting (solid) and non-bursting (hatched) in infant (light red) and juvenile (dark red) SAM-LB reared pups. Infant control-reared vs juvenile control-reared, X^2^=1.55, *p*=0.213; Infant SAM-LB-reared vs juvenile SAM-LB-reared, X^2^=2.77, *p*=0.096; Infant control-reared vs infant SAM-LB-reared, X^2^=12.37, *p*<0.001; Juvenile control-reared vs juvenile SAM-LB-reared, X^2^=0.074, *p*=0.786. All data reported as mean ± SEM. \**p*<0.05, \*\**p*<0.01, \*\*\**p*<0.001, ^#^*p*<0.1

#### Caregiving adversity produces an early hyper-bursting phenotype in infant LHb neurons

We observed qualitative differences in the interspike interval histograms (Extended Data Figure 2B), therefore we further analyzed the features underlying these differences, including burst firing. We categorized cells as bursting (Figure 2G) if they displayed a CV greater than 0.4 and if more than 20% of recorded spikes were fired in a burst ^101^. Although we saw an increase in the number of bursting cells in infants versus juveniles in control-reared pups, this did not reach significance (Figure 2H; X^2^=1.55, *p*=0.213). By contrast, we observed significantly more bursting cells in SAM-LB reared infant pups compared to age-matched controls (Figure 2I; X^2^=12.37, *p*=0.004). Within the same number of active cells, SAM-LB-reared infants showed twice as many bursting cells as controls. Additionally, we saw a large increase in the percent of spikes fired within a burst (% SFB) in infant SAM-LB-reared pups (Extended Data Figure 2C; 2-way ANOVA, main effect of rearing: *F*_(1, 142)_=4.025, *p*=0.047; interaction: *F*_(1, 142)_=8.549, *p*=0.004) and a trend (2D; *p*=0.055) towards longer interburst intervals in infant SAM-LB-reared pups compared to age-matched controls, suggesting these cells are only firing in a bursting pattern with long pauses between each burst. Of note, we saw no differences in the duration of bursts or number of spikes within a burst between any group (Extended Data Figure 2E-F; 2-way ANOVA, *p*>0.05), suggesting the bursting parameters are similar in SAM-LB-reared and control-reared pups. Altogether, these data suggest that ECA disrupts the typical developmental transition of LHb activity by prematurely and significantly increasing bursting.

### Experiment 3

#### Combining social and threat cues engages the developing LHb

The preceding set of experiments characterized the impacts of developmental age and early rearing environment on baseline measures of habenula structure and function. Our next step was to assess these impacts on task-dependent activity of the LHb in infant and juvenile pups.

In adults, the LHb has been shown to be responsive to aversive cues or threats ^25,26^. Therefore, we employed metabolic imaging to assess habenula activity in response to a threatening stimulus (a series of mild shocks) with a social partner present, compared to threat alone (Figure 3A-B). There was a main effect of age (Figure 3C; 3-way ANOVA, *F*_(1, 97)_=2.652, *p*=0.107), rearing (3-way ANOVA, *F*_(1, 97)_=9.170, *p*=0.003), and social context (3-way ANOVA, *F*_(1, 97)_=9.603, *p*=0.003) on 2-DG uptake in the LHb. We also observed an interaction of age and social context (3-way ANOVA, *F*_(1, 97)_=6.632, *p*=0.012). *Post hoc* comparisons revealed significant reductions in 2-DG uptake in the LHb of juvenile control-reared pups compared to infants when shocked in a nonsocial context (*p*=0.016). Additionally, in juvenile control-reared pups, 2-DG uptake was significantly higher in the LHb when shocked in a social context compared to a nonsocial context (*p*=0.008). These effects were blunted in SAM-LB-reared pups (all comparisons, *p*>0.05). We observed a similar pattern across the lateral and medial subregions of the LHb and the MHb (Extended Data Figure 3B-D). These data suggest that the LHb processes social and threat information differently across development.

**Figure 3.**
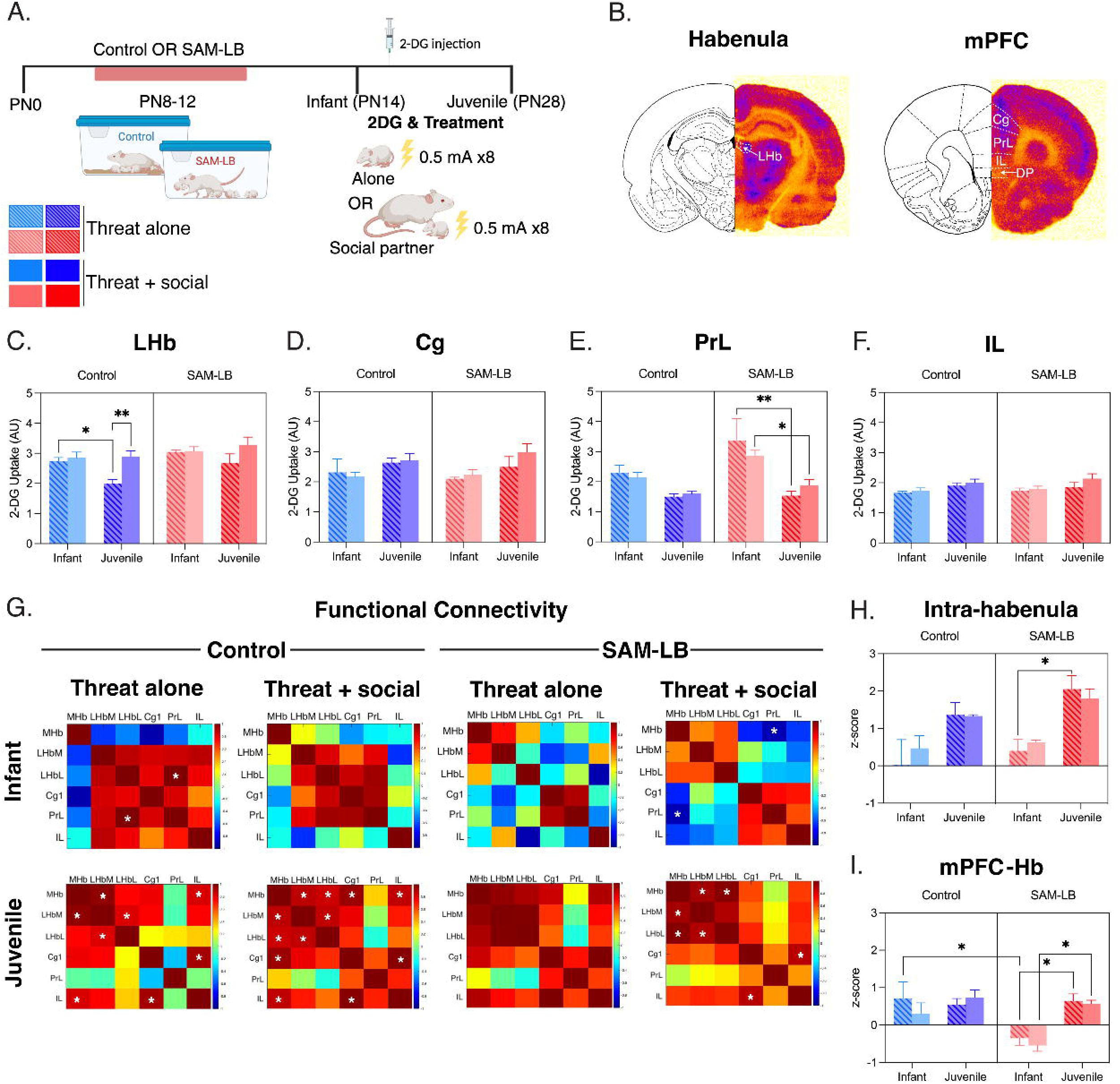
Combining social and threat cues engages the developing LHb. (A) Schematic illustrating experimental timeline. (B) Example autoradiograph showing 2-DG uptake in the LHb (left) and mPFC (right). (C) Average 2-DG uptake in the LHb. (D) Average 2-DG uptake in the Cg. (E) Average 2-DG uptake in the PrL. (F) Average 2-DG uptake in the IL. (G) Functional connectivity heatmaps in infant control (left, top), juvenile control (left, bottom), infant SAM-LB (right, top), and juvenile SAM-LB reared (right, bottom) pups. Red indicates a strong positive correlation. Blue indicates a strong negative correlation. Starred squares reach significance. (H) Average intra-habenula functional connectivity. (I) Average functional connectivity between the mPFC (Cg, PrL, IL) and Hb. All data reported as mean ± SEM. \**p*<0.05, \*\**p*<0.01, \*\*\**p*<0.001, ^#^*p*<0.1

In adults, the mPFC sends a glutamatergic projection to the LHb which conveys stress information ^42^ and inhibits social approach behavior ^40^. Furthermore, the late-developing mPFC is a locus of dysfunction following ECA ^19,102–104^. Therefore, we sought to determine whether this region showed any involvement in processing of social and threat information. We looked across the mPFC at the infralimbic (IL), prelimbic (PrL), and cingulate (Cg1) cortex. We observed a main effect of age in the Cg1 (Figure 3D; 3-way ANOVA, *F*_(1, 33)_=7.772, *p*=0.009), PrL (Figure 3E; 3-way ANOVA, *F*_(1, 33)_= 31.63, *p*<0.0001), and IL (Figure 3F; 3-way ANOVA, *F*_(1, 33)_=6.786, *p*=0.0134). We observed a main effect of rearing specifically in the PrL (3-way ANOVA, *F*_(1, 33)_=8.239, *p*=0.007). The IL and Cg1 both showed a similar pattern, with increased 2-DG uptake with age in control-reared and SAM-LB-reared pups, although the *post hoc* comparisons failed to reach significance (all comparisons, *p*>0.05). By contrast, in the PrL, we observed reduced 2-DG uptake with age in SAM-LB-reared pups regardless of social context (social: *p*=0.031; nonsocial; *p*=0.002). We observed a similar trend in control-reared pups, but it did not reach significance (all comparisons, *p*>0.05). We observed no effects of age or rearing on 2-DG uptake in other cortical regions (Extended Data Figure 3). Altogether, these data suggest that the subregions of the mPFC convey threat information differently across development and, importantly, that threat information is processed by these regions regardless of social context. These data also demonstrate that ECA alters how the PrL subregion of the mPFC processes threat information in both infant and juvenile pups.

We next assessed functional connectivity within the LHb by generating bivariate correlation matrices (Figure 3G). We observed that correlated activity within habenula subnuclei increased across development (Figure 3H; main effect of age, *F*_(1, 16)_=25.92, *p*=0.0001). These effects reached statistical significance in SAM-LB-reared pups shocked alone (*p*=0.048) but not controls (p=0.168). By contrast, we found that age, rearing, and social context interacted to impact functional connectivity between the mPFC and LHb (Figure 3I). We observed a main effect of age (*F*_(1,208)_=17.69, *p*<0.001), a main effect of rearing (*F*_(1, 64)_=8.762, *p*=0.004) and an interaction between age and rearing (*F*_(1, 64)_=7.617, *p*=0.008). *Post hoc* pairwise comparisons revealed a developmental increase in mPFC-LHb connectivity in SAM-LB-reared pups shocked in both social (*p*=0.017) and nonsocial (*p*=0.049) contexts. Interestingly, SAM-LB-reared pups showed significantly reduced mPFC-LHb connectivity in infancy compared to controls (*p*=0.027) but reached comparable levels of connectivity by the juvenile period (*p*>0.05).

These effects are recapitulated when assessing correlated activity between specific pairs of brain regions across conditions (Table 2). These results also show that intra-habenular connectivity (e.g. LHbL→MHb) is impacted by developmental age, regardless of early care quality. By contrast, adversity, developmental age, and social context all impact connectivity between the LHb and PrL. Together, these data indicate the combination of social and threatening cues differentially engages the infant versus juvenile LHb. Importantly, the mPFC-LHb circuit appears uniquely vulnerable to ECA, suggesting potentially distinct roles for the LHb and mPFC-LHb projections in developing social behavior.

**Table 2.**
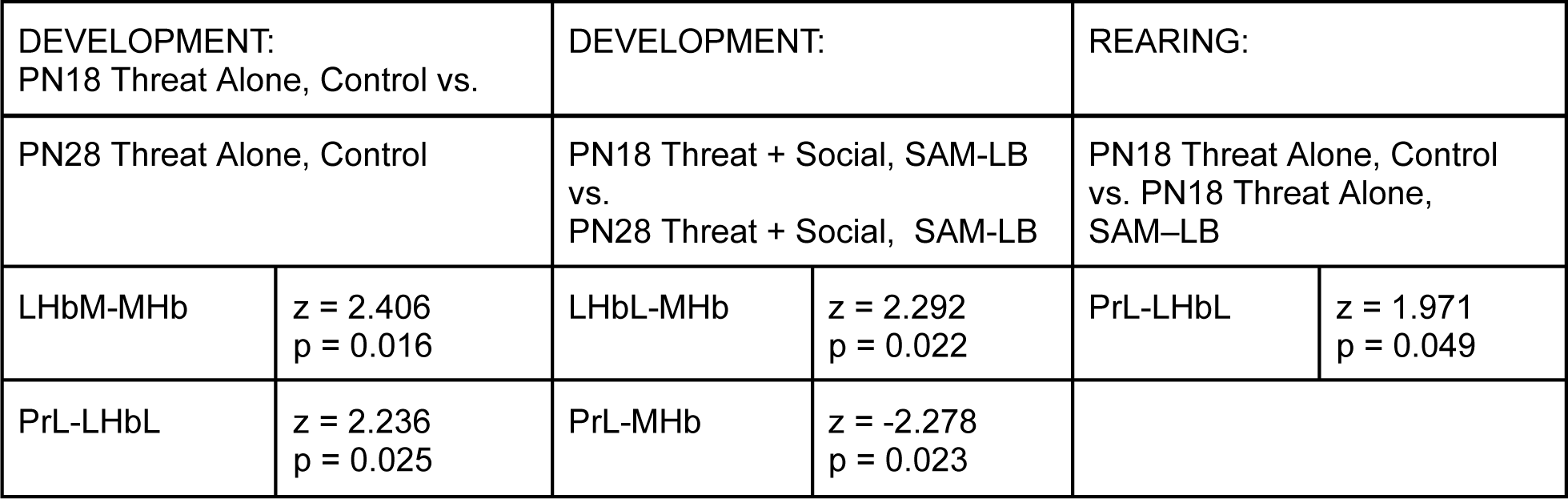
Significant Differences in Pairwise Correlations for ROIs. Abbreviations: LHbL, lateral subdivision of lateral habenula; LHbM, medial subdivision of the lateral habenula; MHb, medial habenula; Cg1, cingulate cortex; PrL, prelimbic cortex.

### Experiment 4

#### LHb activity plays developmentally distinct roles in social approach behavior under threat

The correlational data shown in Figure 3 suggests that how the LHb processes social and threat information changes across development. Therefore, we next asked how the LHb controls the behavioral response to social and threat cues. To causally test this, we used chemogenetics to inhibit the LHb during social behavior testing with and without threat (Figure 4A-B). In this experiment, we used ambient predator odor as a threat cue instead of a tail shock so that social behavior and locomotion would remain undisturbed during the behavior test. Previous work has shown that infant rats demonstrate behavioral and neural responses to natural predator odor^72,105,106^.

**Figure 4.**
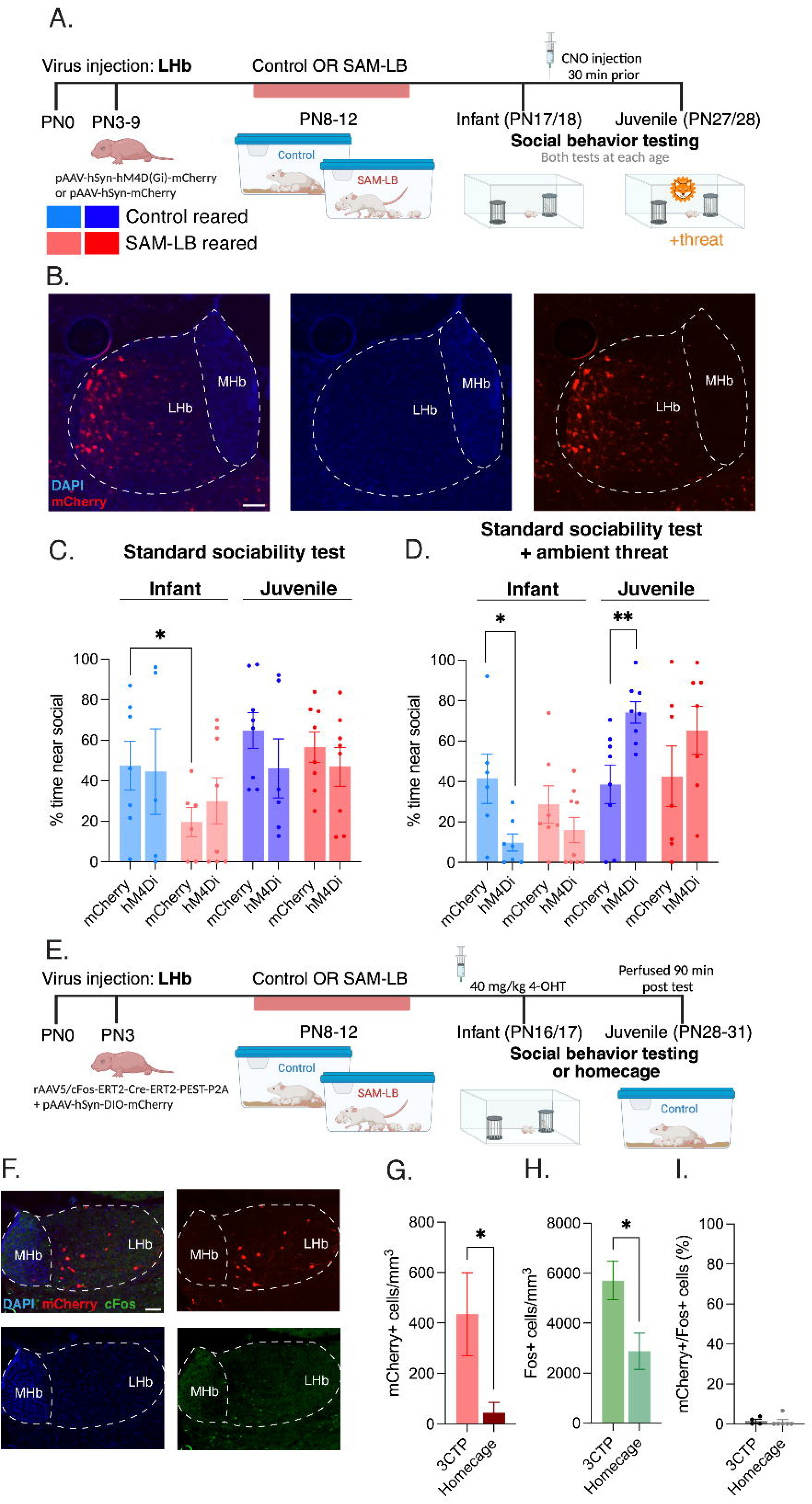
LHb activity plays developmentally distinct roles in social approach behavior under threat. (A) Schematic illustrating experimental timeline. (B) Example histology of DREADD (mCherry) in the LHb. (C) Average time near social partner in infant control (light blue), juvenile control (dark blue), infant SAM-LB (light red), and juvenile SAM-LB (dark red). (D) Average time near social partner when ambient threat present. (E) Schematic illustrating experimental timeline. (F) Example histology of tagged (mCherry) and Fos+ (GFP) cells in the LHb. (G) Average mCherry+ cells in the LHb following sociability testing or homecage. (H) Average Fos+ cells in the LHb following sociability testing or homecage. (I) Overlap between mCherry+ and Fos+ cells (mCherry+/Fos+) in the LHb following sociability testing or homecage. *Post hoc* comparisons performed between hM4Di and mCherry control groups. All data reported as mean ± SEM. Scale bar = 100 um.\**p*<0.05, \*\**p*<0.01, \*\*\**p*<0.001, ^#^*p*<0.1

Consistent with previous results ^7,8^, infant SAM-LB-reared pups showed decreased social approach in a standard sociability test (Figure 4C; *t*_(11)_=1.884, *p*=0.043). We observed a main effect of age on social approach behavior in the standard 3CT (3-way ANOVA, *F*_(1,48)_=5.034, *p*=0.030). *Post hoc* comparisons revealed CNO administration to pups transduced with the inhibitory DREADD in LHb principal neurons had no effect on social approach behavior in the standard 3CT in infant (all comparisons, *p*>0.05) pups, suggesting the LHb is not involved in this behavior at these early ages.

The results of Experiment 3 (Figure 3) suggested that the combined stimuli of threat and social context specifically engages the LHb. Therefore, we added a threat context to the sociability test by introducing ambient predator odor and suppressing LHb activity at each age. We observed a main effect of age (3-way ANOVA, *F*_(1,51)_=21.37, *p*<0.001) and an interaction of age with the inhibitory DREADD in LHb principal neurons on social behavior (3-way ANOVA, *F*_(1,51)_=14.48; *p*<0.001). Remarkably, *post hoc* comparisons revealed that, in infant control-reared pups, inhibition of the LHb suppressed social approach when threat was present (Figure 4D; *p*=0.032). By contrast, in juvenile control-reared pups, this manipulation *increased* social approach when threat was present (Figure 4D; *p*=0.008). These patterns were observed regardless of rearing but did not reach significance in SAM-LB-reared pups (all comparisons, *p*>0.05). Notably, we did not see an effect of the inhibitory DREADD during the habituation period in infants or juveniles of either group (Extended Data Figure 4; all comparisons, *p*>0.05). Together, these data suggest the LHb serves as a neural substrate supporting the transition from early, approach-biased social behavior to a more adult-like balance of approach and avoidance as individuals mature and encounter threats.

We next sought to determine whether overlapping or distinct ensembles of LHb neurons are involved in these developmentally distinct roles of the LHb. To accomplish this, we adapted a viral Fos-CreERT system for pups to longitudinally label active neurons during a sociability test under threat in infancy and as juveniles (Figure 4E-F). This method permitted us to identify ensembles active during the infant test (mCherry-tagged) and ensembles active during the juvenile test (Fos-tagged). Importantly, we included a control group that spent time in their homecage at the two ages instead of undergoing the sociability test with ambient threat. We found that there were significantly more tagged mCherry+ cells (Figure 4G; *p*=0.023) and Fos+ cells (Figure 4H; *p*=0.032) in the LHb following social behavior under threat as compared to homecage controls, suggesting the combined threat-social exposure preferentially engaged the LHb. Importantly, we saw almost no (<1%) overlap between mCherry and Fos cells in the LHb (Figure 4I), suggesting the developmentally distinct roles of the LHb in social approach behavior under threat are mediated by distinct neural ensembles.

### Experiment 5

#### mPFC projection to the LHb drives social behavior deficits in infant pups following early caregiving adversity

The results of Experiments 1-3 show robust impacts of adversity rearing on the LHb. However, broad chemogenetic suppression of LHb activity did not ameliorate social behavior deficits following ECA, suggesting the need for a more specific microcircuit manipulation. In adults, the mPFC sends a unidirectional glutamatergic projection to the LHb that conveys stress information ^42^ and suppresses social approach behavior ^40^. Given the adversity effects on mPFC activity and mPFC-LHb connectivity (Figure 3), we next used an intersectional strategy to target LHb-projecting mPFC neurons (Figure 5A-B). To our knowledge, this the first implementation of this preparation in young, awake-behaving rat pups. We observed a main effect of age (3-way ANOVA, *F*_(1,45)_=10.04, *p*=0.003) and an interaction of age and the inhibitory DREADD (3-way ANOVA, *F*_(1,45)_=5.565, *p*=0.023). *Post hoc* comparisons revealed that suppressing this projection rescued typical social behavior in adversity-reared infants (Figure 5C; *p*=0.006). Again, we did not see an effect of the inhibitory DREADD during the habituation period in infants or juveniles of either group (Extended Data Figure 5; all comparisons *p*>0.05). We saw no effect of suppressing this projection in control-reared infants (Figure 5C; *p*=0.448) or in juveniles of either group (Figure 5C; all comparisons, *p*>0.05). Additionally, we only observed an effect of suppressing the mPFC projection to the LHb in the standard sociability test and *not* when threat was present (Figure 5D; all comparisons, *p*>0.05), suggesting that this projection does not play a role in social behavior under threat. Rather, these data suggest that abnormal mPFC-LHb activity drives social behavior deficits in infant pups following ECA.

**Figure 5.**
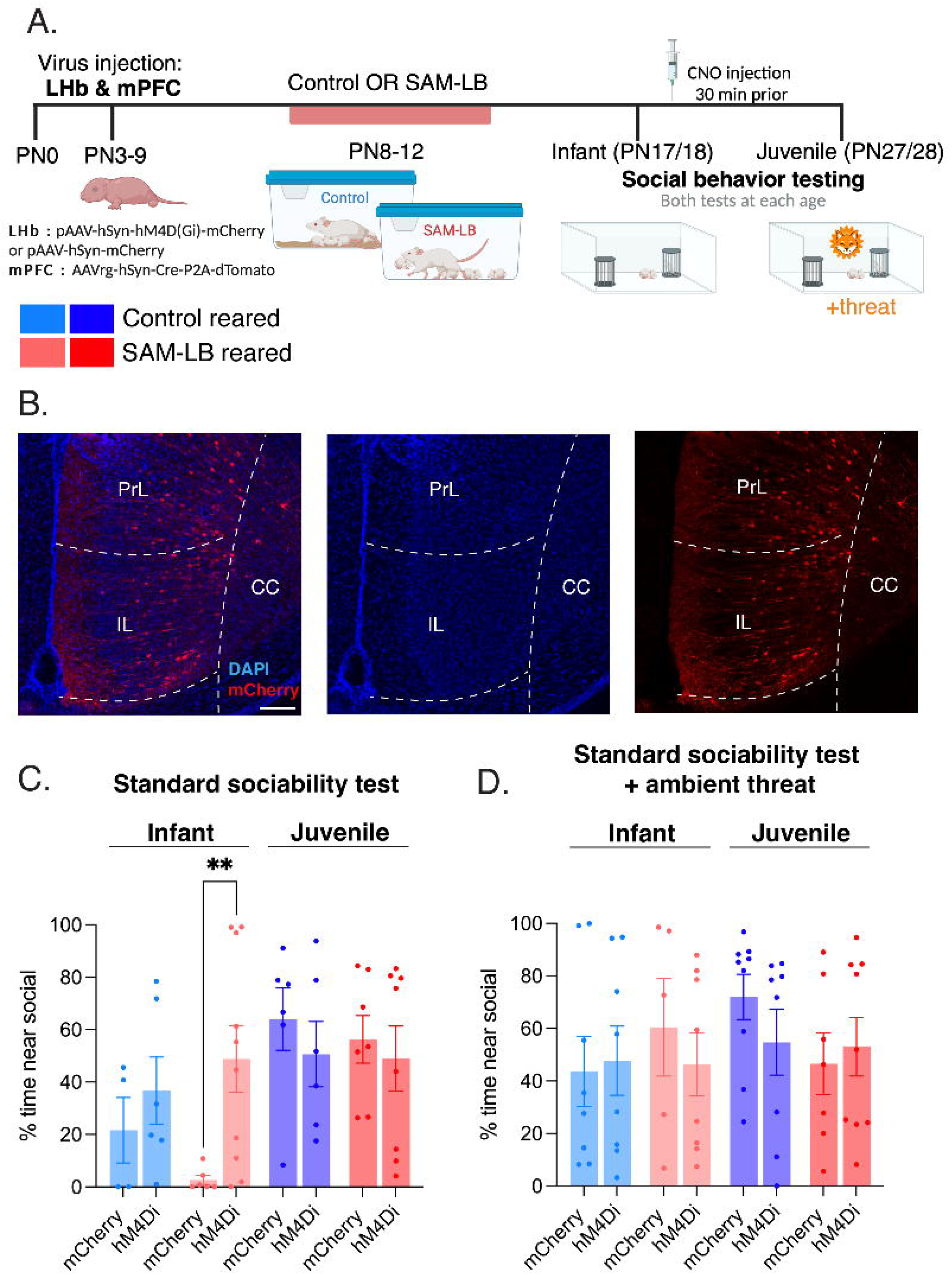
mPFC projection to LHb drives social behavior deficits in infant pups following early caregiving adversity. (A) Schematic illustrating experimental timeline. (B) Example histology of DREADD (mCherry) into the mPFC. (C) Average time near social partner in infant control (light blue), juvenile control (dark blue), infant SAM-LB (light red), and juvenile SAM-LB (dark red). Post hoc comparisons performed between hM4Di and mCherry control groups. (D) Average time near social partner when ambient threat present. *Post hoc* comparisons performed between hM4Di and mCherry control groups. All data reported as mean ± SEM. Scale bar = 100 um.\**p*<0.05, \*\**p*<0.01, \*\*\**p*<0.001, ^#^*p*<0.1

## Discussion

Flexible social behavior is crucial throughout development, but particularly during developmental transitions when environmental demands are in flux. Furthermore, these transitional periods, such as from infancy to the juvenile stage, are marked by heightened vulnerability to environmental impacts. However, neither the circuit mechanisms promoting typical social flexibility nor its perturbation are well understood. In adults, the LHb has been robustly linked with behavioral flexibility and highlighted as a locus of dysfunction in psychopathology ^25,27–31^. However, little is known about its typical functional development or its vulnerability to early adversity. Here, we combined structural and functional assessments to demonstrate how the LHb integrates social and threat cues to promote social behavior flexibility in early development, and how this is perturbed with ECA to impair social behavior.

We assessed the structural and functional ontogeny of the LHb using measures of volume, protein expression, spontaneous activity, metabolic imaging, and loss-of-function chemogenetic circuit dissection in infant and juvenile rats. We demonstrated that across infant to juvenile development, there is an increase in habenular volume and changes in expression of excitatory and inhibitory proteins that correspond to a developmental increase in the spontaneous firing rate of LHb neurons. We next demonstrated developmentally-distinct roles of the LHb in social behavior under threat, with the LHb promoting social approach under threat in infants but *inhibiting* social approach under threat in juveniles. By contrast, we found that ECA inhibits growth of the LHb and alters neuromodulatory protein expression. This corresponded to increased bursting activity in the LHb in infant rat pups that experienced ECA. Importantly, we observed altered connectivity between the LHb and the mPFC in animals that experienced ECA, and subsequently found that social behavior deficits can be rescued by inhibiting this projection. Altogether, these data suggest that ECA produces immediate structural and functional changes to the infant habenular circuit and identifies the LHb as an early locus of dysfunction in social behavior.

### Functional changes to habenular activity across development

LHb neurons express a variety of excitatory, inhibitory, and neuromodulatory receptors that influence their activity. We observed a developmental upregulation of CaMKIIβ and downregulation of GABA_B_ receptors. We saw a similar trend following ECA. CaMKIIβ is expressed in excitatory neurons and plays a key role in activity-dependent plasticity ^107^, while GABA_B_ receptor activity mediates slow, prolonged neuronal inhibition ^108^, thus these changes suggest a developmental increase in LHb excitability. Consistent with this, we found a developmental increase in spontaneous, or tonic, firing of LHb neurons. Interestingly, we saw no effects of ECA on this developmental change in tonic firing, consistent with previous work investigating cerebral blood flow in the LHb in human patients with MDD which found no differences in resting state cerebral blood flow, a measure of tonic activity ^87^.

The LHb also receives numerous neuromodulatory inputs from the ventral tegmental area, dorsal raphe nucleus, lateral hypothalamus, and more ^41,93–96^. Expression of the corresponding neuromodulatory receptors in the LHb are associated with anxiety-like and depressive-like behavior in rodents ^39,97,98,109^. We found developmental decreases to 5HT2C receptors under control conditions but not following ECA. Additionally, we saw a reduction in ESR1 expression from infancy to the juvenile period following ECA. Previous work in adult rodents has shown that intra-LHb administration of ESR agonists suppressed spontaneous firing and c-Fos expression ^98,110^ and that intra-LHb administration of 5HT2C agonists increased spontaneous firing ^109,111^. Importantly, these treatments also reduced anxiety-like and depressive-like behaviors ^98,110^.

Among the most robust adversity effects that we observed was an increase in the proportion of bursting cells and an increase in the proportion of spikes fired within bursts in infant rats. In fact, although we observed the same number of spontaneously active cells in control-reared and SAM-LB-reared infant rat pups, we observed twice as many bursting cells in SAM-LB-reared pups as compared to controls. These results demonstrate how ECA produces acute changes in phasic activity of LHb neurons. Such an increase in phasic activity is consistent with studies done in human patients with MDD, which found abnormal phasic responses to aversive cues in the habenula ^87^. Adversity-induced habenular bursting may also alter phasic signaling in downstream monoaminergic targets. If phasic responses in the LHb and midbrain are disrupted, processing of salient cues, which relies on phasic responses to aversive and rewarding stimuli, will likely be impaired ^20,112–115^. Another interpretation of these results, showing decreased firing outside of bursts, is more redundancy in the LHb coding signal. This parallels increased cross-frequency cortical oscillatory coupling we previously observed immediately following ECA ^70^. Such impacts to long-range information flow may be one mechanism by which ECA impacts the ability to make accurate and timely decisions in scenarios requiring assessment of salient cues across the lifespan, including complex social settings.

Importantly, it is notable that we see these changes to excitability very soon after the ECA experience, which highlights the importance of early detection of biomarkers of stress. By the juvenile period, these effects appear to normalize, which is consistent with previous work showing latent periods preceding emergence of adult psychopathology following infant adversity ^116–118^. The mechanisms underlying this effect remain unknown, but compensatory changes in ESR1 and 5HT2RC expression observed here (Figure 1F-G) are a potential substrate.

### Perturbed habenular growth curve as an early biomarker for psychopathology

Changes to habenular volume are correlated with a variety of psychiatric conditions in humans. Reduced habenular volume has been shown in adult patients with schizophrenia ^86^, bipolar disorder ^50^, and MDD ^50,87^. By contrast, pediatric patients with autism spectrum disorder have been shown to have significantly larger habenular volumes ^119^. We found that developmental increases in habenular volume were blunted following ECA, and that this phenotype emerges at very young ages. Therefore, habenular volume could be used as an early biomarker for development of later psychopathology and for subsequent early intervention. However, the mechanism producing reduced habenular volume remains unclear. Reduced LHb volume may be consistent with our electrophysiology data, which shows fewer active cells in the LHb of juvenile pups that experienced ECA. Future work will be needed to elucidate this mechanism, however, excitotoxicity due to large increases in burst firing, as we observed following ECA, is one possible target. Indeed, reduced neuronal cell counts are observed in the LHb in adult psychopathology ^49^.

### The lateral habenula integrates developmental and contextual demands to promote flexible behavior

In adults, the LHb has been identified as a functional locus for behavioral flexibility in tasks with changing reward or punishment contingencies ^29–31^. Our results using chemogenetic suppression of the LHb demonstrate that this brain region also encodes developmental demands to promote behavioral transitions. Specifically, inhibiting the LHb produced opposite effects before and after weaning, with LHb suppression decreasing social approach in infants and increasing social approach in juveniles when a threat is present. This behavioral transition is consistent with a transition in ecological niche favoring infant approach towards a caregiver when threatened, versus a more avoidant strategy as the individual matures and faces social configurations that may themselves be threatening ^8–10^. Previously, we have demonstrated that a dopaminergic projection to the basolateral amygdala inhibits social approach as infants mature into weaning age ^8^. Here, we show that the LHb, upstream of this circuitry, integrates threat and social cues to coordinate this developmental transition when threat is present. To identify a potential neural substrate for this change, we adapted a viral activity-dependent tagging strategy for developing rats, which showed that distinct neural ensembles are activated by the social-threat test in infants versus juveniles. Future work will be necessary to more fully characterize the neural mechanisms supporting this change across weaning, although targeting downstream, functionally-dissociated projections, such as LHb-VTA-NAc and LHb-VTA-mPFC, is a promising approach ^120,121^.

### A corticohabenular projection inhibits social approach after adversity

Although broad chemogenetic suppression of LHb neurons identified age-specific functions for the region, these effects were specific to a task involving combined threat and social cues. Furthermore, they were observed regardless of early caregiving quality. However, our initial experiments showed robust impacts of early adversity on LHb structure (volume, protein expression) and function (burst firing, task-dependent metabolic mapping), in addition to replicating previous work showing impaired social approach behavior ^7,8^. To reconcile these findings, we implemented the first intersectional viral strategy for chemogenetics in awake, behaving rat pups to specifically target glutamatergic LHb-projecting mPFC neurons. Our metabolic imaging assay highlighted the mPFC-LHb interface as a locus of integration of development, social context, and early adversity. Furthermore, this projection suppresses social approach behavior in adults ^40^. We found that inhibiting these neurons rescued social approach in adversity-reared infants. Thus, these results functionally identify the mPFC-LHb microcircuit as a substrate for adversity-induced social deficits. Notably, the PrL subregion of the mPFC may be critically important in driving adversity-induced social deficits. Increased activity of the PrL has been shown to increase fear generalization in adult rats ^122–124^ and humans with post-traumatic stress disorder ^125,126^. We observed increased activity in the PrL following ECA, suggesting adversity may impair fear memory discrimination leading to social deficits. This is supported by human literature showing that fear generalization is correlated with early adversity in children and adolescents ^127,128^.

### Conclusion

In summary, the current results highlight the LHb as an early locus of social behavior flexibility as well as a structural and functional target of ECA. Using chemogenetics adapted for infant rat pups, we observed that this brain region promotes social approach under threat in infants, while inhibiting this behavior in juveniles. Activity-dependent tagging showed that this behavior was supported by distinct neural ensembles at each age. Furthermore, we found that ECA perturbed habenula growth curves along measures of volume, neuromodulatory protein expression, intrinsic activity, and functional connectivity with the mPFC. Using an intersectional chemogenetic strategy, we identified a specific population of LHb-projecting mPFC neurons that inhibits social approach following early adversity.

The major significance of our work is that it provides evidence for the LHb as an early substrate for social behavior flexibility and a specific microcircuit that translates early adversity to infant social deficits. Additionally, our results connect ECA with known circuit mechanisms of psychopathology and identify immediate biomarkers of adversity within the LHb. Altogether, these findings demonstrate a clinically-relevant target for therapeutic interventions following early trauma and uncover clues about the developmental role of the LHb in social behavior.

## Acknowledgements

The authors would like to thank the following funding sources: NIH BRAIN Initiative R00MH124434, R01MH133456, and BBRF NARSAD Young Investigator Award to MO. Schematics created using BioRender.com.

