## Extended Data for "The lateral habenula integrates age and experience to promote social transitions in developing rats"

**Extended Data Figure 1. Original uncropped western blots used for preparation of Figure 1.** (A) CaMKIIβ western blots with tubulin controls. (B) $GABA_B$ western blots with tubulin controls. (C) 5HTR2C western blots with tubulin controls. (D) ESR1 western blots with tubulin controls.

**Extended Data Figure 2. Detailed assessment of bursting parameters.** (A) Schematic illustrating experimental timeline. (B) Average interspike interval histograms for control infants (light blue), control juveniles (dark blue), SAM-LB infants (pink), and SAM-LB juveniles (red). (C) Average percent spikes fired in bursts. (D) Average interburst interval. (E) Average burst duration. (F) Average number of spikes per burst. All data reported as mean ± SEM. $^{*}p<0.05$, $^{**}p<0.01$, $^{***}p<0.001$, $^{\#}p<0.1$

**Extended Data Figure 3. Expanded assessment of ROIs using 2-DG metabolic mapping** (A) Schematic illustrating experimental timeline. (B) Average 2-DG uptake in the MHb. (C) Average 2-DG uptake in the LHbM. (D) Average 2-DG uptake in the LHbL. (E-M) Average 2-DG uptake in cortical subregions: anterior insular cortex (AIV), lateral occipital cortex (LO), claustrum (CL), dysgranular insular cortex (DI), medial occipital cortex (MO), dorsolateral occipital cortex (DLO), granular insular cortex (GI), ventrolateral occipital cortex (VO), and dorsal agranular insular cortex (AID). All data reported as mean ± SEM. $^{*}p<0.05$, $^{**}p<0.01$, $^{***}p<0.001$, $^{\#}p<0.1$

**Extended Data Figure 4. Distance traveled during habituation and social behavior testing in Experiment 4.** (A) Schematic illustrating experimental timeline of LHb DREADD experiment. (B) Average distance traveled during standard sociability test habituation in infant control (light blue), juvenile control (dark blue), infant SAM-LB (light red), and juvenile SAM-LB (dark red). (C) Average distance traveled during habituation when ambient threat present. (D) Average distance traveled during standard sociability test. (E) Average distance traveled during sociability test when ambient threat present. *Post hoc* comparisons performed between hM4Di and mCherry control groups. All data reported as mean ± SEM. $^{*}p<0.05$, $^{**}p<0.01$, $^{***}p<0.001$, $^{\#}p<0.1$

**Extended Data Figure 5. Distance traveled during habituation and social behavior testing in Experiment 5.** (A) Schematic illustrating experimental timeline of mPFC-LHb DREADD experiment. (B) Average distance traveled during standard sociability test habituation in infant control (light blue), juvenile control (dark blue), infant SAM-LB (light red), and juvenile SAM-LB (dark red). (C) Average distance traveled during habituation when ambient threat present. (D) Average distance traveled during standard sociability test. (E) Average distance traveled during sociability test when ambient threat present. *Post hoc* comparisons performed between hM4Di and mCherry control groups. All data reported as mean ± SEM. $^{*}p<0.05$, $^{**}p<0.01$, $^{***}p<0.001$, $^{\#}p<0.1$

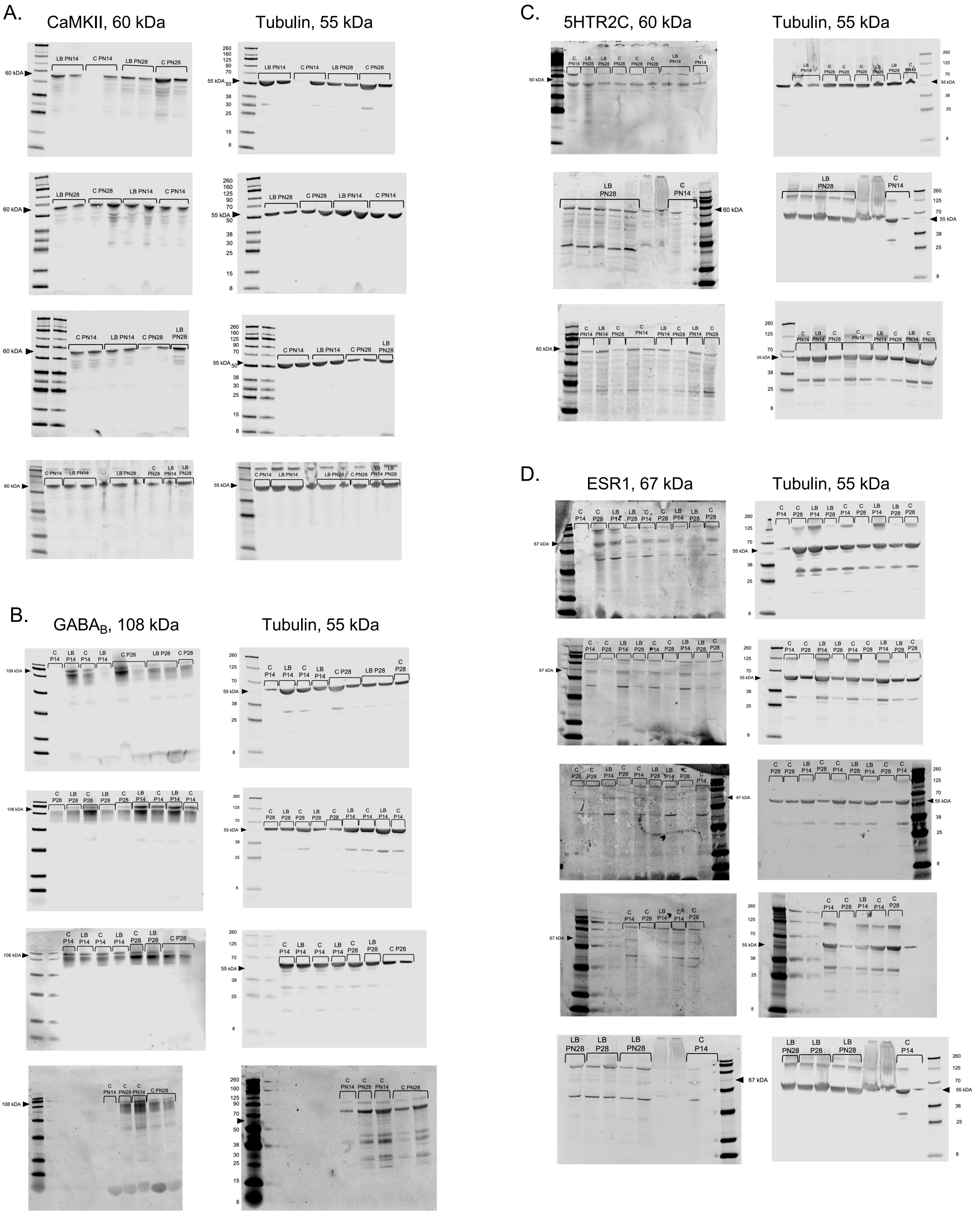
A.
CaMKII, 60 kDa
Tubulin, 55 kDa
B.
GABA$_B$, 108 kDa
Tubulin, 55 kDa
C.
5HTR2C, 60 kDa
Tubulin, 55 kDa
D.
ESR1, 67 kDa
Tubulin, 55 kDa

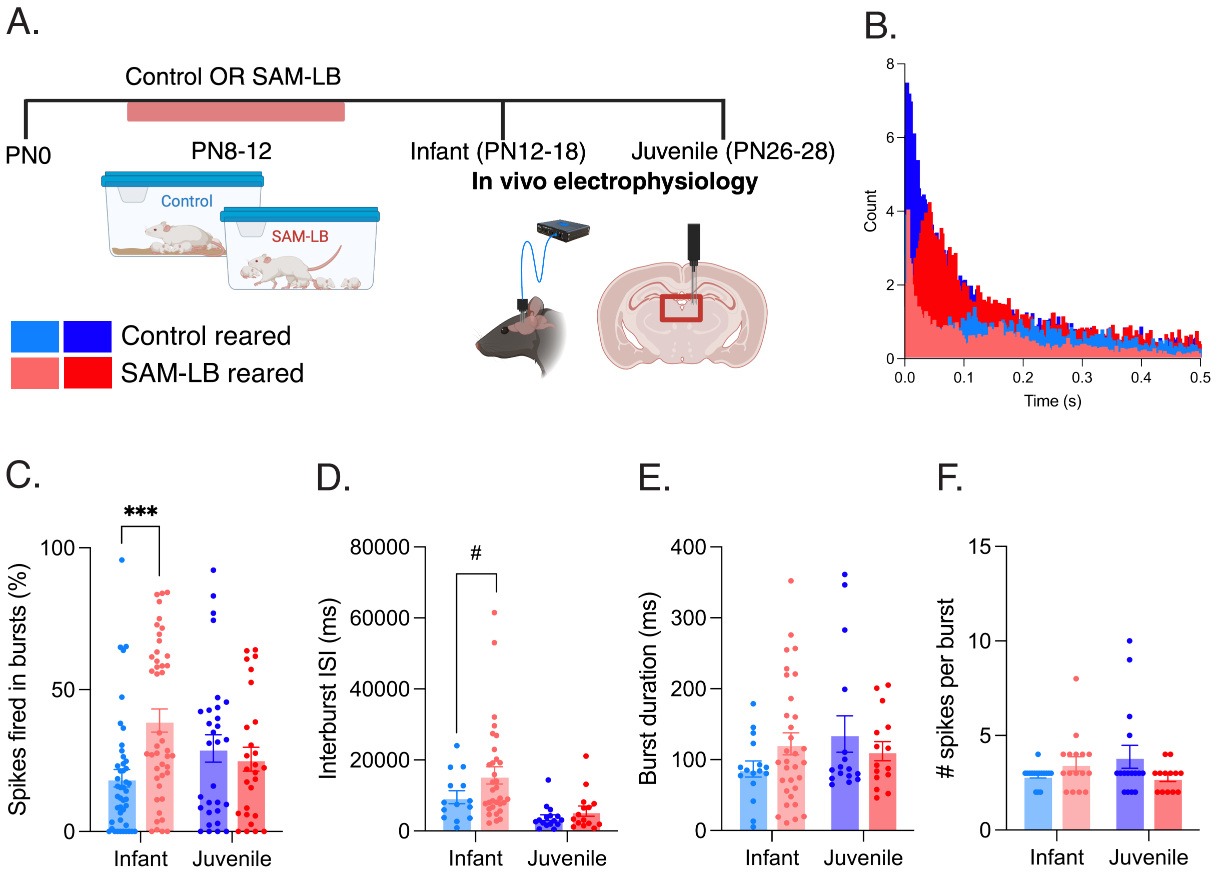

A.
Control OR SAM-LB
PN0
PN8-12
Infant (PN12-18)
Juvenile (PN26-28)
In vivo electrophysiology
Control
SAM-LB
Control reared
SAM-LB reared
B.
Count
Time (s)
C.
Spikes fired in bursts (%)
***
Infant
Juvenile
D.
Interburst ISI (ms)
#
Infant
Juvenile
E.
Burst duration (ms)
Infant
Juvenile
F.
### spikes per burst
Infant
Juvenile

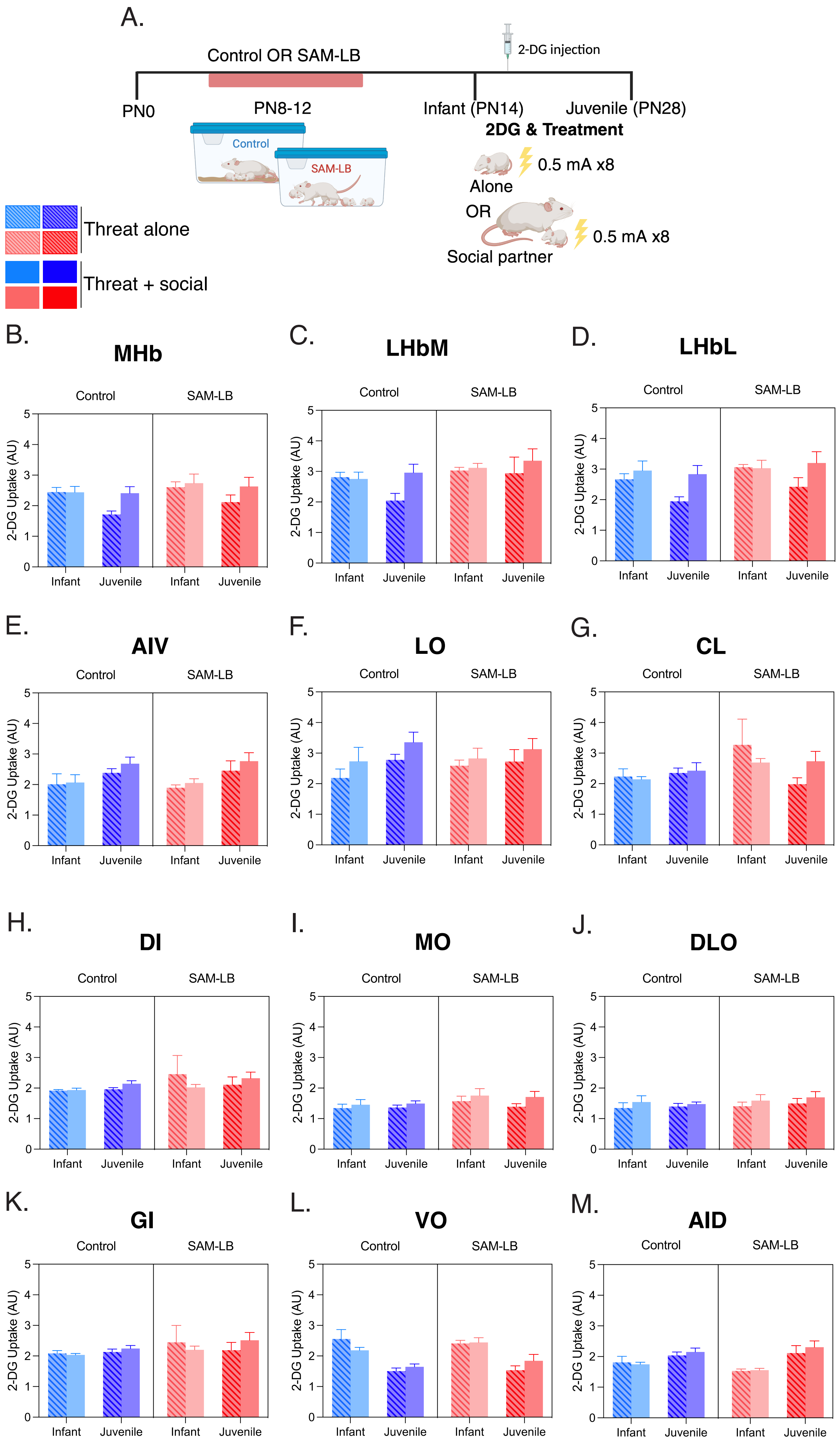

A.
Control OR SAM-LB
2-DG injection
PN0
PN8-12
Infant (PN14)
Juvenile (PN28)
2DG & Treatment
Control
SAM-LB
0.5 mA x8
Alone
OR
0.5 mA x8
Social partner
Threat alone
Threat + social
B.
MHb
C.
LHbM
D.
LHbL
E.
AIV
F.
LO
G.
CL
H.
DI
I.
MO
J.
DLO
K.
GI
L.
VO
M.
AID
Control
SAM-LB
2-DG Uptake (AU)
Infant
Juvenile

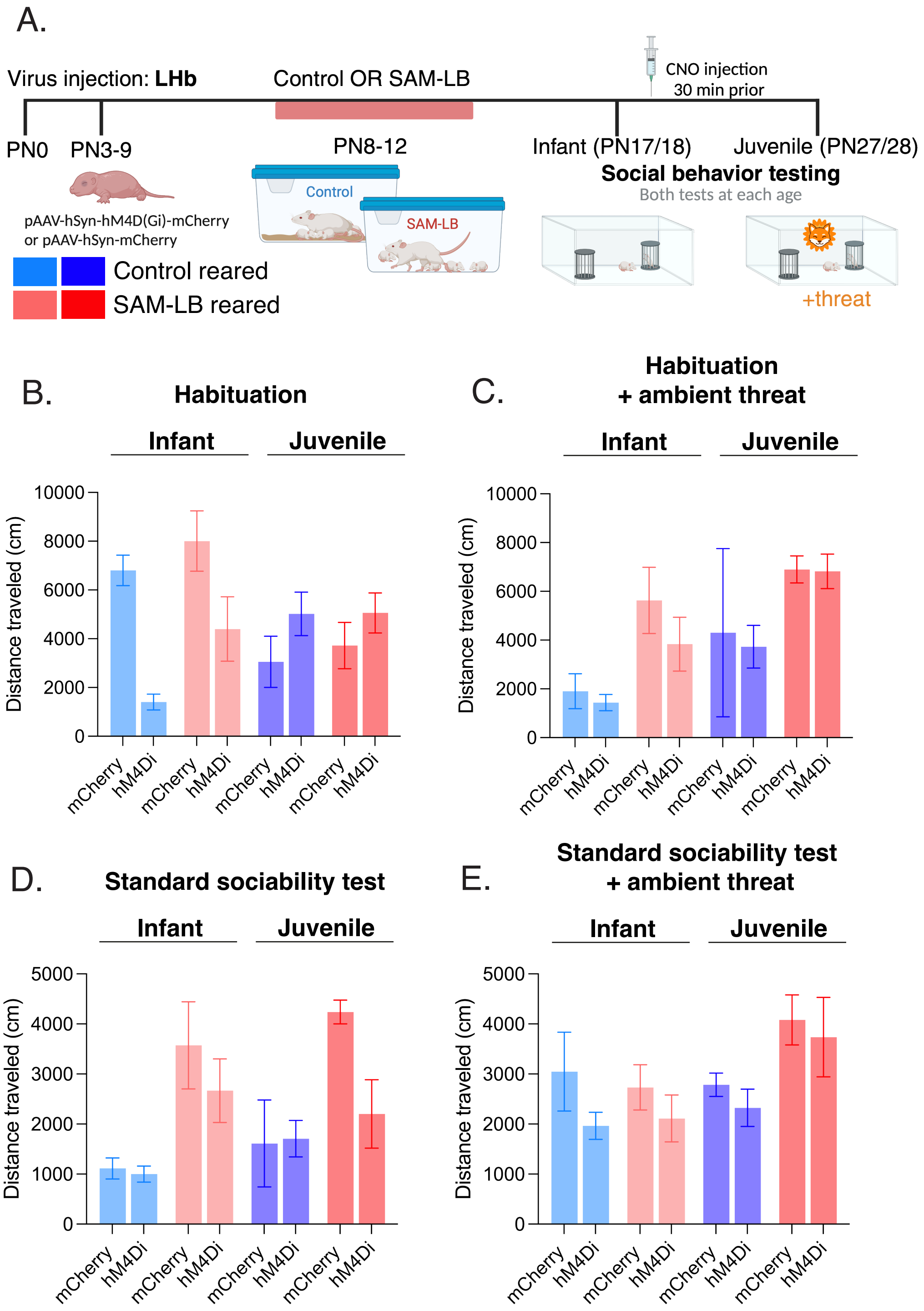
A.
Virus injection: LHb
Control OR SAM-LB
CNO injection
30 min prior
PN0
PN3-9
PN8-12
Infant (PN17/18)
Juvenile (PN27/28)
Social behavior testing
Both tests at each age
pAAV-hSyn-hM4D(Gi)-mCherry
or pAAV-hSyn-mCherry
Control
SAM-LB
Control reared
SAM-LB reared
+threat
B.
Habituation
C.
Habituation
+ ambient threat
D.
Standard sociability test
E.
Standard sociability test
+ ambient threat
Infant
Juvenile
Distance traveled (cm)
mCherry
hM4Di

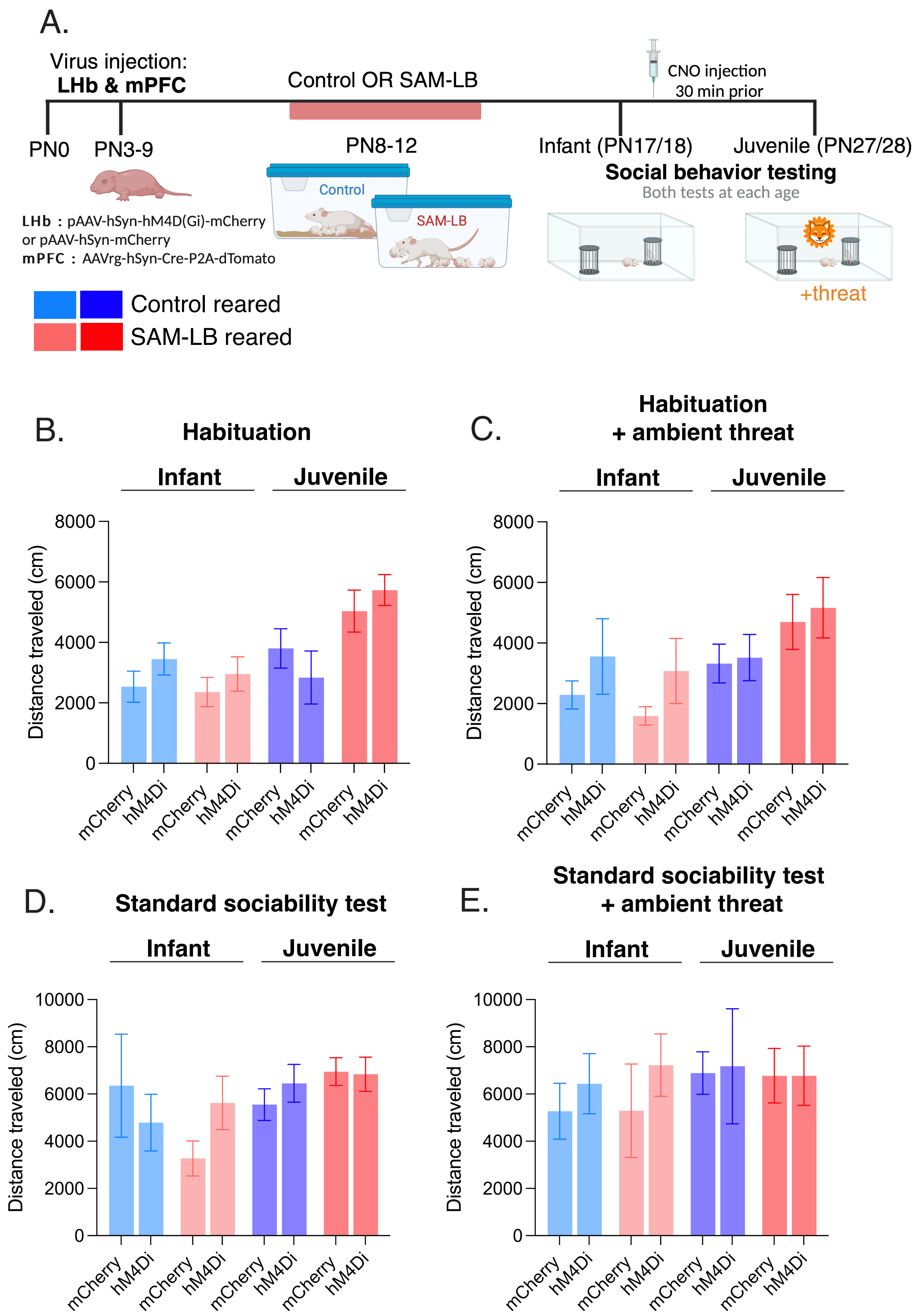
A.
Virus injection:
LHb & mPFC
Control OR SAM-LB
CNO injection
30 min prior
PN0
PN3-9
PN8-12
Infant (PN17/18)
Juvenile (PN27/28)
Social behavior testing
Both tests at each age
Control
SAM-LB
LHb : pAAV-hSyn-hM4D(Gi)-mCherry
or pAAV-hSyn-mCherry
mPFC : AAVrg-hSyn-Cre-P2A-dTomato
+threat
Control reared
SAM-LB reared
B.
Habituation
Infant
Juvenile
Distance traveled (cm)
mCherry
hM4Di
C.
Habituation
+ ambient threat
D.
Standard sociability test
E.
Standard sociability test
+ ambient threat
